# Statistical inference of virus evolution using ancestral recombination graphs

**DOI:** 10.64898/2026.10.02.756209

**Authors:** Christopher H.C. Leung, Charlotte J. Houldcroft, Aylwyn Scally

## Abstract

Ancestral recombination graphs (ARGs) represent the coalescent history at every locus in the genome, subject to mutation and recombination. Modelling viruses using the coalescent with recombination remains less explored, due to the potential for complex, multi-scale dynamics. Using perturbative methods, we establish mathematical bounds that describe the balance between inter-host and within-host coalescence, and demonstrate that the highly-recombinant Epstein-Barr Virus (EBV) coalesces predominantly at the inter-host level. We then modify SINGER, a tool from diploid genetics, for haploid ARG inference with fine-grained variable recombination rates. We provide insight into the paradox of high recombination rates and slow linkage disequilibrium decay, as ARG-based methods potentially allow us to disentangle epistatic effects from shared ancestry, and illustrate this using the coalescent topologies of EBV’s host cell-entry glycoproteins. Our findings demonstrate that ARG-based methods recover gene-level selective structure, and provide a general method for analysing virus phylodynamics in the limit of small quantities.

## Introduction

Understanding virus evolution is important for the analysis of viral function and selective constraints, as well as for the tracking and prediction of outbreaks. Viral evolution is often modelled as a coalescent process, which traces sequences back in time to their common ancestors and produces a tree topology. However, reticulate gene transfer between lineages can occur during recombination. Consequently, each position in the genome may have its own coalescent tree; at recombination breakpoints, a subset of the topology changes. The combination of these marginal trees gives the ancestral recombination graph (ARG).

Two generative models underpin ARG inference, distinguished by the direction in which the graph is built. The coalescent with recombination (Hudson, 1983) works backwards in time: lineages coalesce as in the standard coalescent, but also split at recombination events, allowing a single lineage to carry ancestral material from two parents. The sequentially Markovian coalescent (SMC) approximation of the ARG (Wiuf and Hein, 1999) instead moves along the genome. Starting from the marginal tree at one end, recombination breakpoints prune and re-graft subtrees, and the correlated sequence of marginal trees forms the graph. Probabilistic models (Rasmussen et al., 2014) or heuristic reconstruction (Minichiello and Durbin, 2006) are the two main paradigms used for ARG inference. These methods have found applications in mapping ancient admixture and introgression in primates and hominids (Tung and Barreiro, 2017; Cousins et al., 2025), inferring selection from genomic data (Hejase et al., 2022), and discovering complex trait associations (Zhu et al., 2025).

Parsimonious ancestral recombination graph reconstructions have been shown to be valuable in describing the reassortment of influenza virus segments (Barrat-Charlaix et al., 2022) and to clarify the evolutionary relationships of recombinant SARS-CoV-2 lineages in large phylogenies (Ignatieva et al., 2021, 2022) – both RNA viruses. Acute RNA viruses have genealogies which are effectively single-scale in their coalescence. Deep latency in a large DNA virus interposes a slow within-host process between transmission events, creating a multi-scale coalescent – a potential obstacle ARG inference has to overcome. Therefore, we introduce probabilistic inference of viral ARGs using a modified version of SINGER, a Bayesian ARG inference method which scales to hundreds of sample sequences (Deng et al., 2025).

Epstein-Barr virus (EBV; *Lymphocryptovirus humangamma4*) is an ubiquitous human gamma herpesvirus, contributing to hundreds of thousands of cancer cases and deaths every year (Wong et al., 2022). EBV has a double-stranded DNA genome of approximately 172kbp and a distinctive pattern of geographic variation: type 1 EBV, which has a B cell tropism, is found worldwide, while type 2, which has a T cell tropism (Coleman et al., 2015), is more commonly detected in Africa (Neves et al., 2017).

EBV is asymptomatically present in 95% of humans, and is primarily transferred via saliva and bodily fluids, with no environmental reservoir. Active EBV shedding primarily occurs in an epithelial, lytic phase, which couples to a B-cell latent reservoir occasionally undergoing lytic reactivation (Zaremba et al., 2025). During latency in B-cells or oropharyngeal epithelial cells, the EBV genome circularises via terminal repeats and is maintained as an episome by the host replicative machinery in the nucleus (Young et al., 2016).

Intra-strain recombination in co-infected host cells may be an enabling mechanism for immune evasion and proliferation. Other studies have inferred a long history of EBV recombination (Berenstein et al., 2018; Agwati et al., 2022), while sub-genomic studies have shown that immune compromised hosts can harbour multiple EBV strains in the same tissue, with recombination inferred to take place (Walling et al., 1992).

EBV is an ideal test of ARG-based methods due to its high recombination rate (Zanella et al., 2019; Agwati et al., 2022), which limits the genomic scope of tree-based analyses. EBV’s annotated immunogenic loci have elevated LD and co-adaptive blocks that retain population structure (Wegner et al., 2019), alongside contradictory topologies recovered from different variant sites. In a freely recombining genome, LD should decay rapidly: shared ancestry is one possible explanation for slow LD decay. If recombination is geographically or structurally restricted, co-inherited allelic combinations are inferred as linked blocks with shared coalescent history. An alternative explanation is epistatic selection, which maintains allelic combinations at functionally coupled loci even as recombination erodes their shared genealogy. Epistatic effects have been previously noted for RNA viruses including vesicular stomatitis virus, SARS-CoV-2 and HIV (Michalakis and Roze, 2004); and DNA viruses herpes simplex virus 1 (an alphaherpesvirus) (Lee et al., 2016) and human cytomegalovirus (a betaherpesvirus) (Peterson et al., 2025). Both mechanisms can sustain LD against the homogenising effect of recombination. By inferring the genetic genealogy at each locus, ARG-based methods could recover evolutionary history and functional genomic constraints with greater power and at a higher resolution than tree-based approaches.

Given that current, at-scale ARG inference tools do not support multi-scale coalescence models such as Dearlove and Wilson (2013); Volz et al. (2017), we first provide theoretical justification that EBV coalescence is a single-scale process due to the transmission bottleneck between individuals. We use perturbation theory to show that two parameters of EBV – viral latency and the separation between reactivation and coalescence timescales – enable single-scale ARG inference. More than an EBV-specific justification, this supplies a general, parameter-based criterion for when a virus admits single-scale ARG inference and when it demands a structured coalescent.

We then extend probabilistic ARG inference to viruses, modifying SINGER to admit haploid genomes and heterogenous recombination rates. We benchmark its performance against ARG simulations matched to EBV’s population parameters. Finally, we infer ARGs from a Type 1 and 2 EBV dataset drawn from Wegner (2017), and use a localised, recombination-aware measure of genetic divergence to analyse the mosaic geographic and disease associations of EBV.

## Results

### EBV coalescence occurs on the inter-host transmission bottleneck scale

The ARG inference methods below assume a single, unstructured coalescent, but in principle the evolution of EBV occurs on two scales. Two coalescence regimes are possible. Prolonged co-infection favours withinhost coalescence: multiple lineages co-occur in immunocompetent carriers throughout infection, as found from analysing latent membrane protein sequences (Sitki-Green et al., 2003). Recombination is inferred to take place in coinfected hosts, and the lineages remain in the host for decades. Deep latency instead favours between-host coalescence: at any moment only a few B-cells are lytically active (Hadinoto et al., 2009), so two lineages are rarely available to coalesce at the same time. To use a single-scale coalescent as a generative model for EBV, we require one of these effects to dominate the other.

The extended latency period limits coalescence to transmission bottlenecks by imposing a severe temporal constraint. If a lineage is active a small fraction *ε* of the time, then two independent lineages are both active only about *ε*^2^ of the time. Therefore, we track the states of a pair of lineages through time and keep terms up to this leading *ε*^2^ coincidence, which bounds the extent of within-host coalescence.

We made this claim precise with perturbation theory, which treats *ε* as a small parameter, and expand the probability of coalescence as a series ordered by the number of reactivations required. This construction is not specific to EBV, and could apply to any virus whose lineages alternate between active and dormant states.

We modelled each viral lineage as a two-state continuous time Markov chain, with rates Lytic 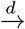 Latent 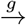 Lytic.

At stationarity it is active with probability

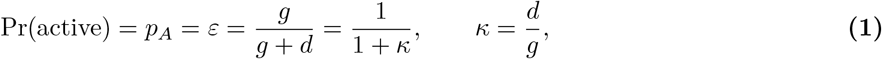

*κ* gives a measure of how strongly latency dominates: deep latency corresponds to large *κ* and small *ε*. To estimate *κ*, we take the average EBV case in the population: an asymptomatic carrier with one in 10, 000 to 100, 000 (Kimura et al., 2008) EBV+ B-cells in a population of 300 billion B-cells (Sender et al., 2023).

Assuming ergodicity and 1-3 lytic B-cells (Hadinoto et al., 2009), this gives *κ* ∈ [1 *×* 10^6^, 3 *×* 10^7^]. This approach is agnostic to the reactivation rate distribution across EBV’s latency programmes.

How does this set a time limit for coalescence? In this Markov chain, because the equilibrium distribution is dominated by the latent state, the relaxation rate (the spectral gap) defines the characteristic time for which a lytic lineage remains active before entering dormancy. This relaxation rate is defined as *R* = *d* + *g*: as *d* is much greater than *g*, transient lytic bursts are mostly isolated from other bursts. Correspondingly, we can reparameterise the transition rates as *g* = *εR* and *d* = (1 − *ε*)*R*.

Now consider the rate of coalescence between two simultaneously active lineages. It occurs at a rate *γ* = 1*/*(*N*_*A*_*τ*_*A*_) per pair of simultaneously lytic B-cell lineages, where *τ*_*A*_ is the viral generation time. Crucially, *N*_*A*_ is the effective number of distinct EBV lineages that are in the lytic compartment, not the total number of virions. This is because when an infected cell bursts, the released virions trace back to their most recent common ancestor within hours. Therefore, coalescence between these virions occurs rapidly relative to the genealogy relating viral samples. Instead, the relevant coalescence occurs when we consider the genetic history between separate lineages that are infecting different lytic foci in the oropharyngeal epithelium.

We now consider a pair of viral lineages. Their joint state (*s*_1_, *s*_2_) admits four possibilities: (Lytic, Lytic), (Lytic, Latent), (Latent, Lytic), and (Latent, Latent). This itself is a continuous-time Markov process, with the doubly-active state (Lytic, Lytic) leading to coalescence at a rate *γ*.

We can describe their joint evolution with a matrix **S**. For EBV, where *d* ≫ *g*, we can take *d* ≈ *R* and *g* = 0, and separate out their fast and slow dynamics into a baseline **S**_0_ and perturbation *ε***S**_1_ respectively. The fast dynamics capture coalescence when the two lineages are both lytic and the rapid lytic-to-latent transition, while the slow matrix *ε***S**_1_ captures the rare reactivations and the *Oε* correction to *d*. **S** is given in Eq. (2) and illustrated in Figure 1.

**Figure 1.**
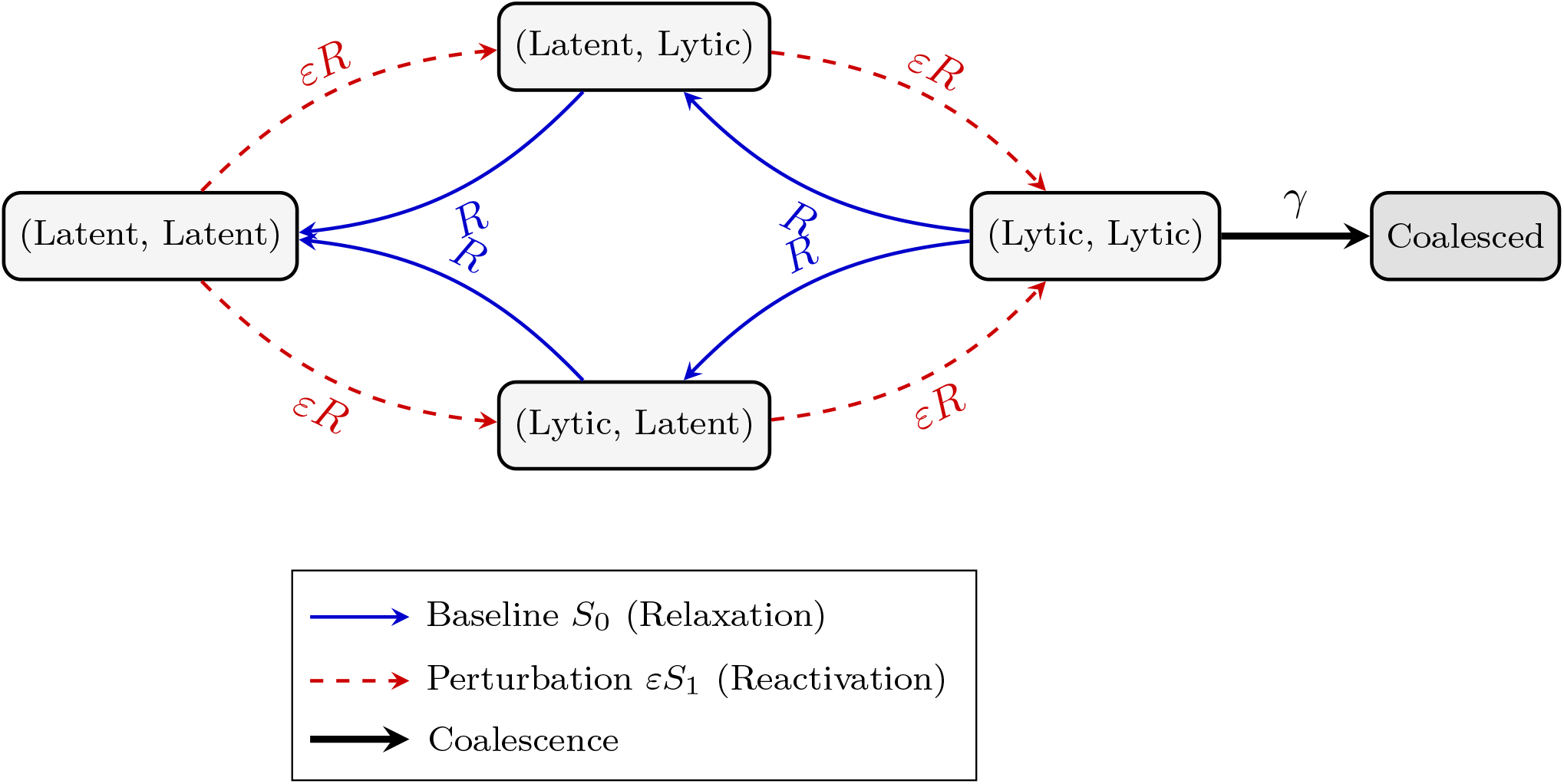
State space and transitions for the joint two-lineage CTMC. The baseline matrix *S*_0_ (solid blue arrows) drives rapid *O*(1) relaxation into the doubly latent state, while the perturbation matrix *εS*_1_ (dashed red arrows) captures rare reactivations into the lytic compartment where coalescence can occur.

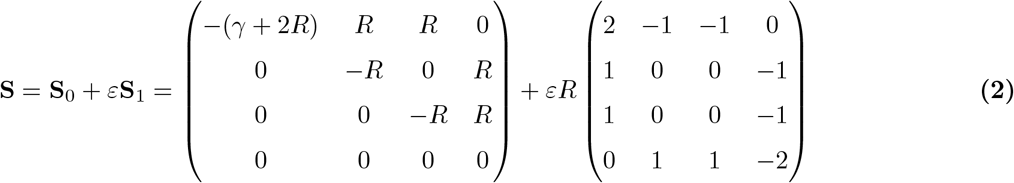

The quantity we seek is the probability that the pair has not coalesced after a co-infection time *T*. This probability is 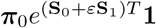. Since *ε* is small, the survival probability can be expanded out as a sum of terms decreasing in size – a convergent Dyson power series (Kato, 1995) – in which each term carries one more reactivation and one more factor of *ε*.

We can keep terms up through *ε* to capture the leading contribution to coalescence by counting reactivating lineages. With none (the zeroth-order term) active, or with one of the pair lytic (the first-order term), coalescence is not possible, so both terms do not contribute. Coalescence first appears with two reactivations, giving to *O*(*ε*^3^)

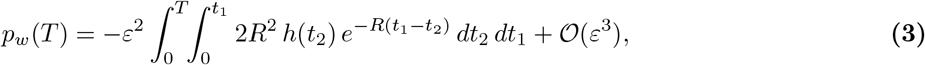

Where 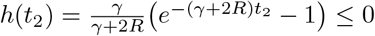.

We can then apply two approximations. When switching is fast compared with coalescence (*R* ≫ *γ*, since lytic bursts terminate quickly while *γ* is held down by the spatial separation of independently reactivated foci) and the co-infection spans many switching times (*RT* ≫ 1, so the brief transients at the ends of the interval are negligible),

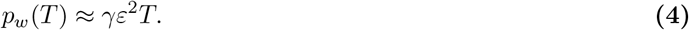

In the case of EBV, this shows that in the limit of rare reactivation, the coalescence rate is bottlenecked by the small fraction *ε*^2^ in which two independent lineages are simultaneously lytic. With *ε*^2^ ∈ [10^−15^, 10^−12^], pairwise within-host coalescence is vanishingly rare. Summing over pairs, a host carrying *N*_*A*_ lineages over a lifetime *T* coalesces internally with probability

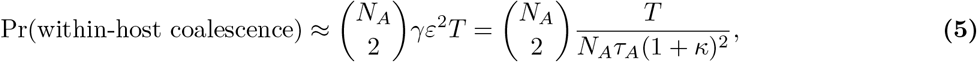

which rises only linearly in *N*_*A*_. Even at *N*_*A*_ = 500, far beyond the 1–3 lineages that drive steady-state shedding, this stays of order 10^−6^ (Figure 2). Within-host coalescence is therefore filtered out at the population scale, and the internal nodes of an ARG collapse within-host variation, summarising only the inter-host transmission bottlenecks.

**Figure 2.**
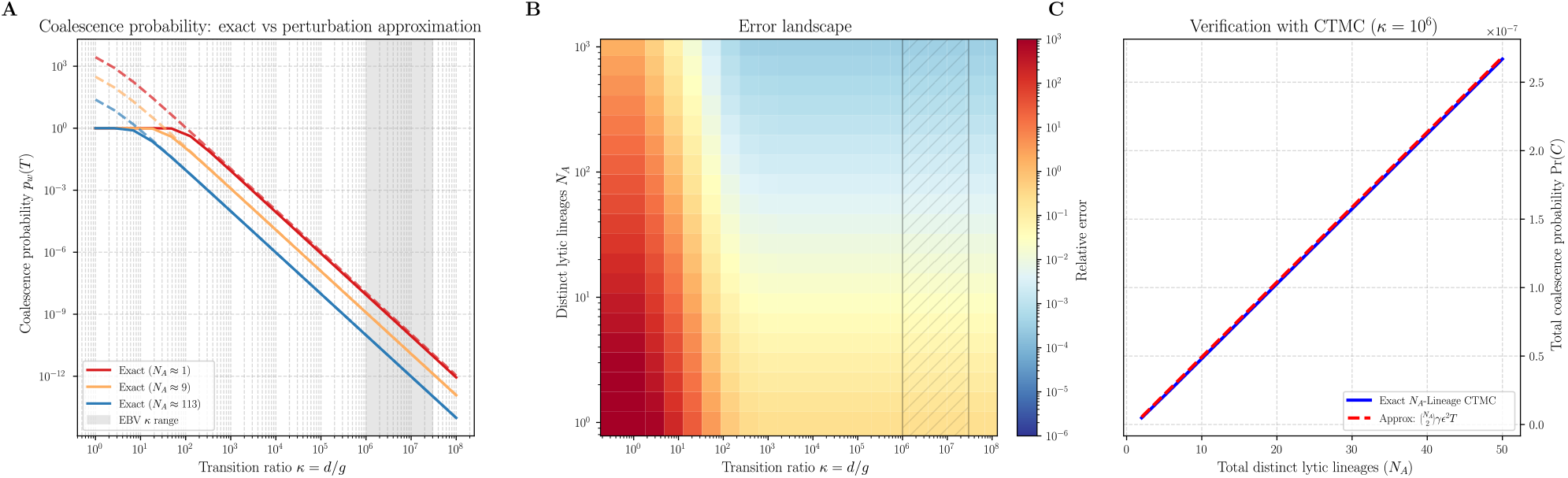
(A) Comparison of pairwise, within-host coalescence probabilities found exactly using mpmath and usinsg the perturbation approximation, with *T* = 60y, *τ* = 2d, *R* = 1d^*−*1^. The approximation breaks down for small *κ* and holds for large *κ*. (B) The absolute error as a fraction of exact pairwise coalescence probability. In the pairwise case, the approximation holds better for larger *N*_*A*_, corresponding to *R* ≫ *γ*. Although EBV has a low effective *N*_*A*_ in most cases, its large *κ* sufficiently suppresses the error. (C) Comparing Eq. (5) against a full simulation of an *N*_*A*_-lineage Markov chain. As *N*_*A*_ increases, the total coalescence probability increases linearly and for *κ* = 10^6^, remains on the 1 in 10 million scale. Picking a large *N*_*A*_ corresponds to a large-scale overlapping reactivation of EBV-infected B-cells, far exceeding the 1-3 lineages that cause steady-state shedding (Hadinoto et al., 2009). Extrapolating, the probability of within-host coalescence with 500 active lineages would still be 1 in a million.

### Recombination is compatible with rare within-host coalescence

We now reconcile this perturbative bound on coalescence with EBV’s frequent recombination. Fundamentally, recombination and coalescence act differently: the marginal genealogy at any single locus is a valid coalescence tree, and recombination simply sets the correlation between neighbouring marginal trees, not the coalescence rate at a given locus. Backwards in a time, a recombination event at a locus reassigns which lineage carries the flanking sequences but leaves the number of lineages ancestral at the locus unchanged. Therefore, it does not change the per-locus coalescence rate described by the model. Moreover, by ignoring recombination, we over-count the probability that a given pair meets at a locus, making the bound conservative.

Recombination in EBV consists of gene conversion and crossover recombination. As gene conversion events are primarily found in the repeat regions, masking SNPs in these regions means crossover recombination dominates. Recombination requires two co-localised genomes during lytic replication in a single cell, whereas coalescence requires two independently reactivated EBV lineages to trace to a common ancestor: these are mechanistically and temporally distinct processes (see Posada et al. (2002)’s comments on recombination and phylogeny estimation). High recombination rates therefore are compatible with our model. Mathematically, this resembles the strong-migration limit of the structured coalescent (Nordborg, 1997).

### Adapting and benchmarking SINGER for viral ARG inference

Therefore, we proceeded with ARG modelling of EBV using a modified fork of SINGER to handle haploid genomes (Supplementary Information). To evaluate SINGER’s performance we benchmarked it alongside RDP5 (Martin et al., 2021), a widely-used program for recombination detection based on a consensus of phylogenetic and sequence-difference signals, in the inference of recombination breakpoints with msprime-simulated multiple sequence alignments (Baumdicker et al., 2022). These were generated using population parameters matched to our EBV dataset, varying the recombination-to-mutation ratio (q=0.02, 0.05, 0.1, 0.2, 0.5). We compared each program’s ability to localise recombination breakpoints to the interval between any two consecutive informative sites (where sequences differ). Additionally for SINGER, we calculated a windowed measure of the time to the most recent common ancestor (TMRCA).

SINGER’s mean inferred values and 95 % credible interval (Figure 3A) closely track the ground truth time for TMRCA and genetic diversity. However, locating breakpoints was substantially more uncertain for both RDP5 and SINGER. At *r/m* = 0.5, across 20 test replicates, SINGER found 1345 of 1959 true intervals and made 1608 false-positive calls, whereas RDP5 found 142 and made 200 false-positive calls. Across the range of *r/m* ratios, SINGER better distinguished recombination-positive from negative inter-SNP intervals than RDP5 (AUROC 0.749-0.779 versus 0.519-0.536; Figure 3B).

**Figure 3.**
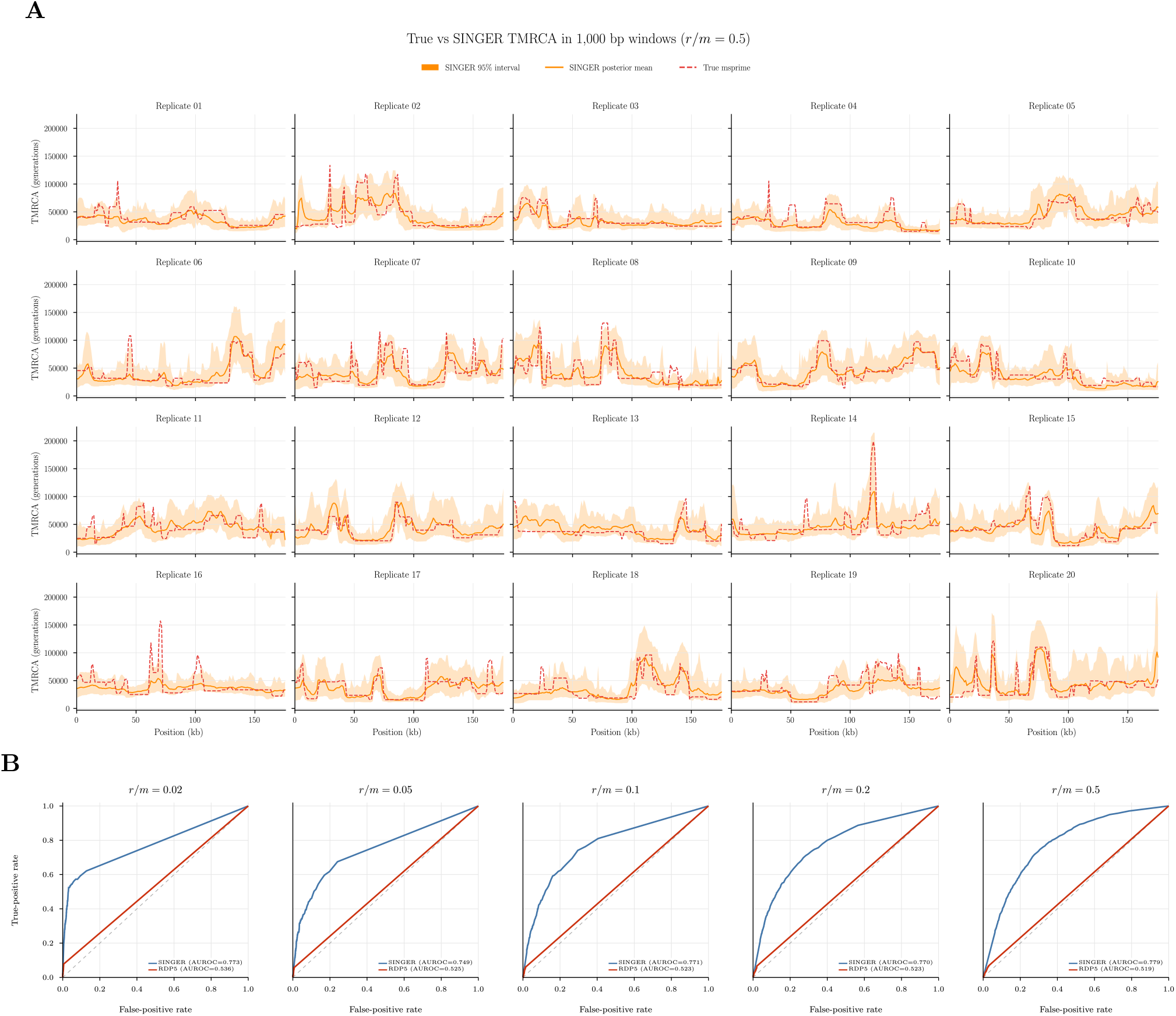
(A) TMRCA in 1000 bp windows across 20 msprime simulation replicates, comparing the true values with SINGER’s posterior mean and 95 % interval. The simulations use population parameters matched to the EBV dataset and *r/m* = 0.5. (B) ROC curves comparing SINGER and RDP5 breakpoint detection across recombination-to-mutation ratios *r/m* = 0.02, 0.05, 0.1, 0.2, 0.5.

Therefore, we sought a measure of coalescent distance that incorporates recombination-induced topology changes, without relying on a strict interpretation of each recombination event. At each marginal tree *T* ^*n*^, the branch-based genetic diversity matrix (Lehmann et al., 2026) 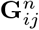 encapsulates the local coalescent distance between sequences *i, j*. Averaging **G** across the marginal trees spanning a genomic interval gives a measure of relatedness that incorporates recombination-induced topology changes.

### Long-range linkage disequilibrium and recombination

We applied SINGER to 147 EBV viral samples (Methods), averaging over 150 ARG samples to infer locus-specific evolutionary relationships and genealogical properties. We also calculated LD between all genomic loci and across the whole sample set. LD between loci separated by large distances along the alignment persists (*p* = 1.1 *×* 10^−3^ (Bonferroni-corrected Fisher’s exact test)) despite pervasive recombination (Figure 4A). Pairwise topological comparison of marginal trees using the Robinson-Foulds metric (Robinson and Foulds, 1981) showed that topological similarity (corresponding to similar ancestral relationships between taxa) is conserved within linkage blocks, but decays over greater distances, and thus does not explain the observed long-range LD (Figure 4B, C). Two possible alternative causes for the allelic correlations represented by LD seen are either geographically-restricted recombination (such that a subset of samples do in fact share ancestry), or long-range epistatic selection maintaining functional haplotypes independently of tree topology. Previous studies have suggested a landscape dominated by purifying selection punctuated by loci exhibiting positive diversification (Santpere et al. (2014); Wegner et al. (2019)).

**Figure 4.**
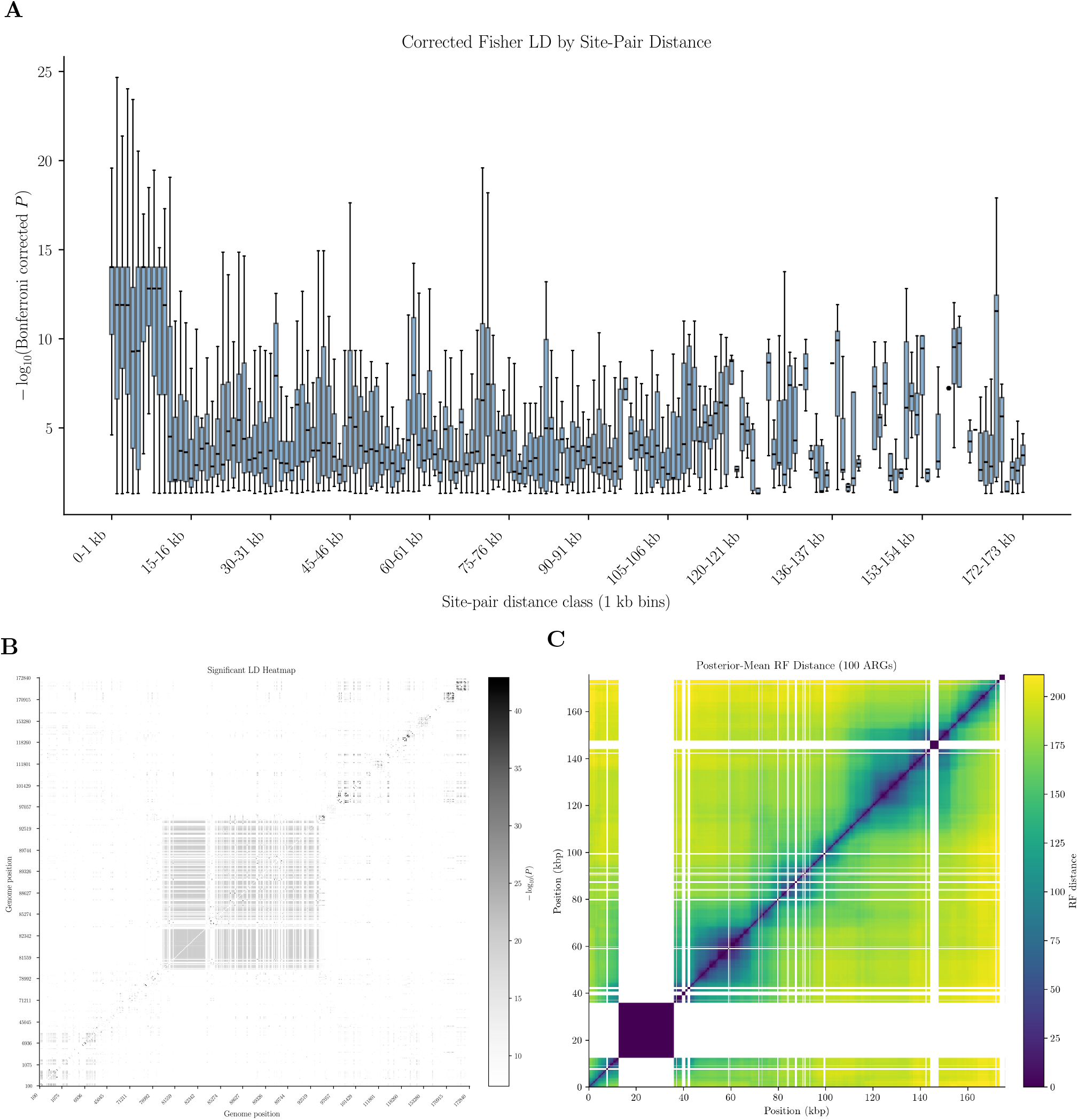
(A) 1kb-binned site-pair distance box-and-whiskers plot of Bonferroni-corrected Fisher test p-values for LD, demonstrating significant allelic associations even at long distances. Our linear distance metric does not account for episomal circularisation. (B) Loci of significant LD along the sequence (note that only significant LD sites are shown). We observe linkage domains of varying intensities in the lytic genes (centre, top-right): considering haplotypes in the context of ORF function and sequence metadata could disentangle functional dependence and ancestral inheritance combined with weak immune selection. (C) The average Robinson-Foulds between each pair of 100bp bins along the genome. White windows correspond to masked repeat regions.

We find that inferred TMRCA varies across the genome (Figure 5 5). This value, and that of locally-inferred *N*_*e*_, is affected by demographic history and population size (which may vary between loci if there is population structure and genomic regions with differing population ancestry), by random sampling of the genealogical process, and by selection. Both purifying and strong positive selection can reduce genetic diversity, and thus inferred TMRCA, while balancing selection can increase it. Inferred ancestry in EBV is characterised by long genomic stretches of low TMRCA, punctuated by short regions of very high TMRCA, inconsistent with neutral variation around a mean value.

**Figure 5.**
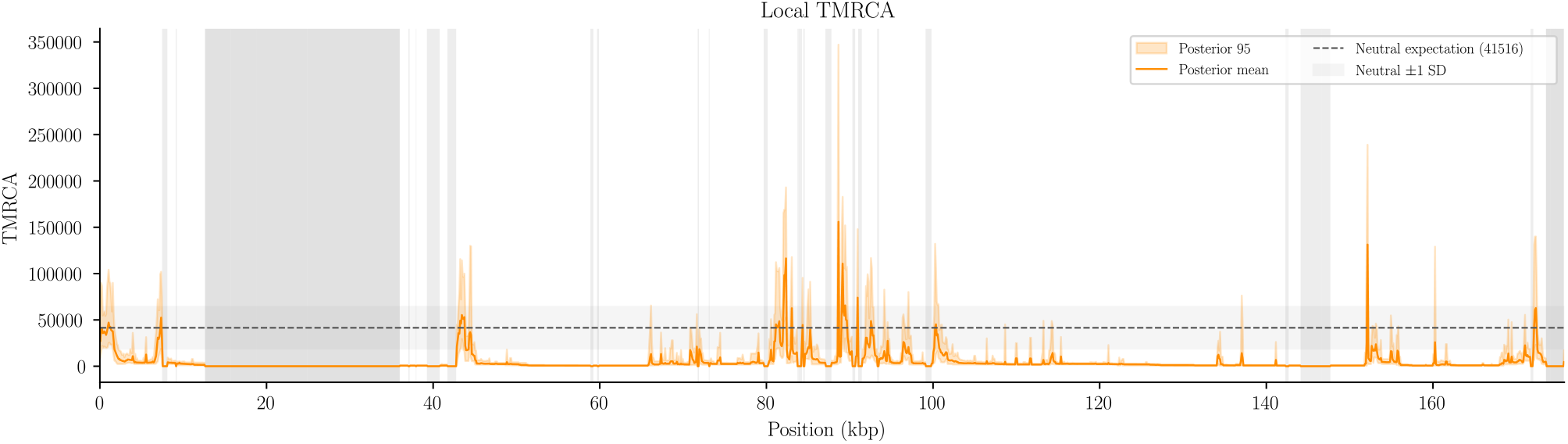
Posterior mean of local TMRCA in 100 bp windows across the EBV genome (orange), with its 95 % posterior interval (shaded), compared against the neutral expected TMRCA (grey, dotted)

### ARG topologies reveal a picture of stratification and host-driven selection: decoupled glycoprotein evolutionary trajectories within EBV’s entry machinery

To investigate the relationship between gene function (and potential functional coupling between genes) and genealogical relatedness as inferred from the ARG, we performed gene-level phylogroup clustering. Applied to the whole genome, the type 1/2 partition is recovered (Blazquez et al., 2025).

As an example of gene-specific analysis based on ARG inference, we focused on the evolutionary relationships of glycoprotein genes within EBV’s entry machinery, *BZLF2, BLLF1* and *EBNA-3B/C*. These show identical clustering (*p* = 7.5 *×* 10^−17^ for Type 2 versus the rest, Fisher’s exact test; Bonferroni threshold *α* = 1.1 *×* 10^−4^). The late gene *BZLF2* encodes glycoprotein gp42, which forms a complex with gL (*BKRF2*) and gH (*BXLF2*) to bind MHC receptors on B-cells and facilitate entry (Li et al., 1995). gL and gH alone are sufficient for epithelial cell entry.

Though all three ORFs encode immunogenic epitopes (Bu et al., 2019), we observe three distinct genealogical regimes across the single functional unit. *BZLF2* follows the Type 1/2 split with a divergence 5.2*×* deeper than the other two ORFs (Table 1) and exhibits purifying selection (*dN/dS* = 0.29). Selection partitions both *BKRF2* and *BXLF2* by geographical region (Figure 6), inter-cluster differentiation is lower, suggesting less-significant, but diversifying selection. *BKRF2* harbours a 7.9-fold greater within-cluster variation than *BXLF2*, which may reflect diversification-constraint asymmetry.

**Table 1.**
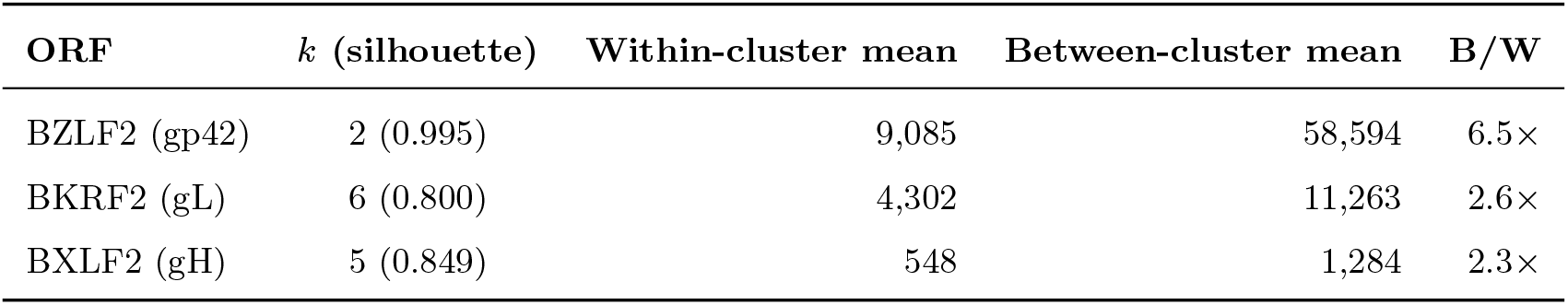
Average posterior mean divergence (in units of generations per bp) within clusters and between clusters for *BZLF2, BKRF2* and *BXLF2*, and the corresponding ratio between the two. *k* is optimised by silhouette score (column 2).

**Table 2.** Parameters and settings for SINGER ARG reference. Zero-coverage windows of the recombination map were floored at 1 % of the genome-wide mean to prevent chain collapse. Also note that the mutation rate used downstream (2.53 × 10^−7^) is the alignment-calibrated value and is more precise than the value passed to the sampler.

| Quantity | Symbol | Value |
| --- | --- | --- |
| *Data* |  |  |
| Alignment | — | $n = 147$ |
| Reference | — | NC_007605 |
| Variant input | — | biallelic SNPs |
| Inference interval | $[s, e]$ | $[0, 175760]$ (MSA coordinates) |
| Repeat masking | $\mathcal{R}$ | enabled (`Repeats_msa.bed`) |
| *Population genetics parameters* |  |  |
| SINGER mutation rate | $m$ | $2.5 \times 10^{-7}$ site$^{-1}$ generation$^{-1}$ |
| SINGER recombination rate | $r$ | $1.25 \times 10^{-7}$ site$^{-1}$ generation$^{-1}$ |
| Recombination-mutation ratio | $r/m$ | 0.5 |
| Recombination map | $r(i)$ | 1 kbp windows |
| Effective population size | $N_e$ | 20 900 (fixed, Watterson) |
| Ploidy | — | 1 |
| Downstream $\mu$ | $\mu$ | $2.53 \times 10^{-7}$ |
| *MCMC* |  |  |
| Retained samples | — | 150 |
| Thinning interval | $t$ | 20 (i.e. 3000 iterations) |
| Burn-in | $b$ | 50 samples (1000 iterations) |
| Posterior window | $[i_{\min}, i_{\max}]$ | $[50, 149]$, giving $K = 100$ |
| Independent chains | — | 1 |

**Figure 6.**
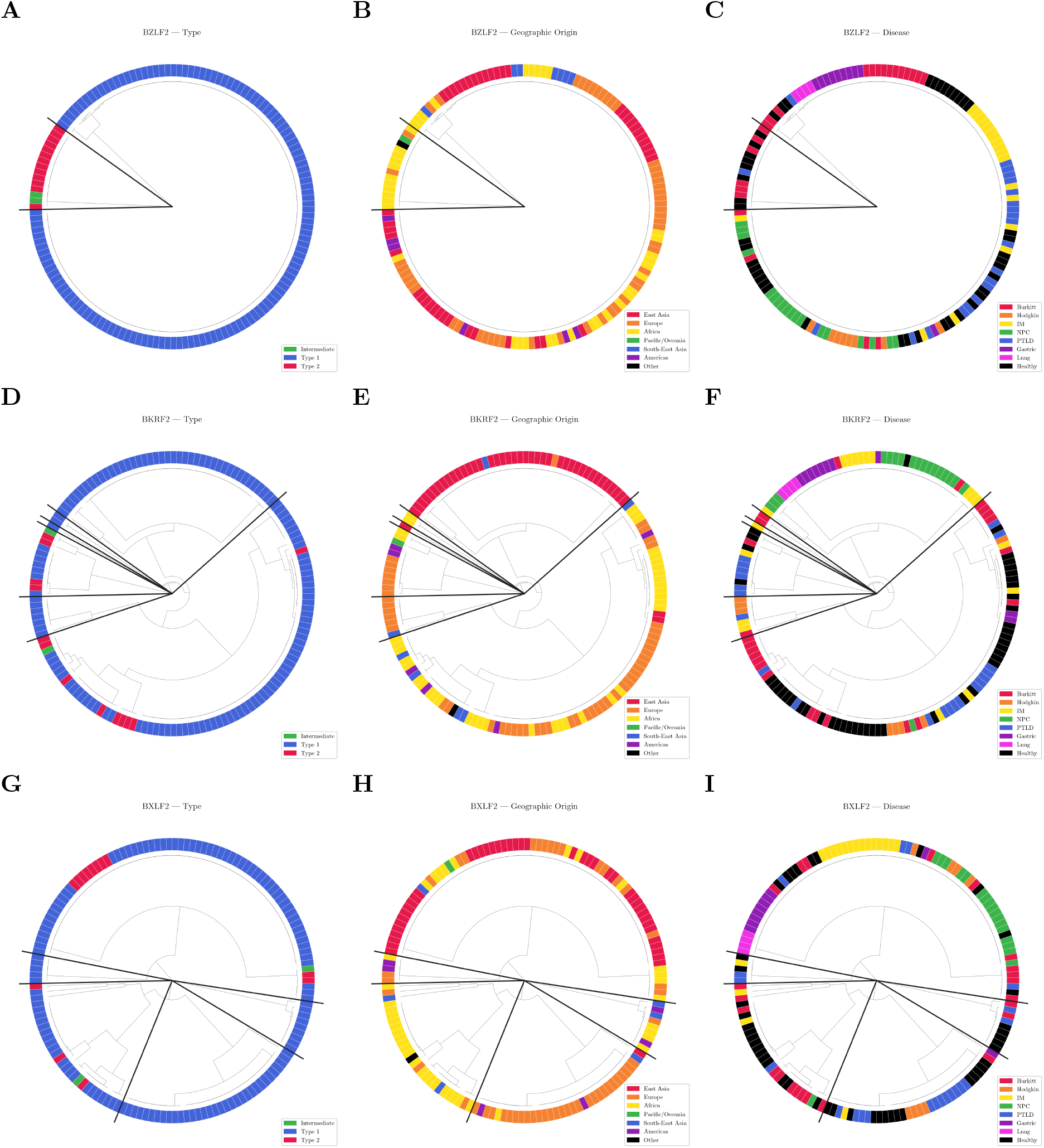
Gene clustering trees for *BZLF2, BKRF2*, and *BXLF2*. Each column is coloured by type, geographical origin, and disease type. Radial lines partition the set into unsupervised clustering groups. Selected co-clustering probabilities (Fisher’s exact test for *k* = 2 clusters, otherwise *χ*^2^ permutation test) [BZLF2 (gp42), BKRF2 (gL), BXLF2 (gH)]: type 1 vs the rest [8.5 *×* 10^*−*21^, 0.010, 0.083], type 2 vs the rest [7.5 *×* 10^*−*17^, 0.045, 0.14], Asia vs the rest [0.0058, 1.0 *×* 10^*−*4^, 1.0 *×* 10^*−*4^]

In light of the transmission-bottleneck result (EBV coalescence occurs on the inter-host transmission bottleneck scale) and the data, we suggest these that the components of a single EBV-surface complex can be subject to overlapping population-level selective and demographic forces. However, by breaking genetic linkage, recombination prevents genetic hitchhiking and retains the distinct coalescent histories of each ORF.

Importantly, geographical clustering is not an inevitable artefact of ARG population structure in non-immunogenic protein-coding genes: for instance, BFLF1, a capsid localisation protein, exhibits within-cluster variation without significant geographic association (*p* = 0.39, permutation test). As an internal negative control, this suggests that geography reflects locus-specific selective pressures, rather than genome-wide demographic structure.

## Discussion

Applying ARG inference to viruses requires a justified coalescent approximation and an assessment of which features of ancestry are recoverable from sequence data alone. We addressed the former with a perturbative description of within-host and between-host coalescence. Simulations of parameter-matched data show that

ARG methods are more sensitive in their detection of recombination breakpoints than existing methods. By integrating over uncertainty in individual recombination histories in EBV, it reveals distinct genealogical histories amongst components of a single molecular complex, allowing the separate analysis of functional coupling, allelic association, and co-inheritance.

### A generalised description of multi-scale coalescence

By modifying the formalism of the seed-bank coalescent (Blath et al., 2016; Cappello et al., 2025) for withinhost dormancy to compute a finite-time pairwise coalescence probability, we showed that the observed history of highly latent viruses is dominated by transmission bottlenecks.

This strengthened our interpretation of SINGER’s inferred topologies as a direct reflection of how recombination affects ancestry over human generations across the genome. Beyond EBV, our method provides a framework for understanding the interplay between within-host and between-host evolution in viruses. The perturbative expansion we introduce depends on the probability of activation (*ε* or *p*_*A*_, and the separation between switching and coalescence timescales, *γ/R*.

Crucially, although our derivation required *ε* ≪ 1 the result of Eq. (4) does not depend on it. As long as *γ/R* ≪ 1, the Markov chain is quasi-stationary. Therefore, by adiabtic elimination the effective rate is *γ*_eff_ = *γε*^2^. An example is HBV, where aggressive, active replication means that *ε* is not small. However, the fast switching between lytic and latent states allows it to satisfy *γ/R* ≪ 1, leading to an ARG that doesn’t collapse fully, but is uniformly scaled by the fraction of the liver in which the virus is actively replicating.

Moreover, our expressions are consistent with the framework of Lythgoe et al. (2017), who propose that the balance between within-host adaptation and transmission determines how much within-host evolution reaches the population. Latent DNA viruses, such as herpesviruses, have small *ε* and *γ/R*, suppressing within-host coalescence and correspondingly to the germline-dominated pole. Where slowly evolving latent lineages are preferentially transmitted. The high degree of latency complements the replication fidelity invoked to explain herpesvirus transmission.

Acute viruses such as influenza and SARS-CoV-2 are at the transmission-dominated evolutionary regime. With a short horizon of *RT* = *O*(1), there are only a few generations before transmission or clearance, giving little time for within-host mutations to accumulate. Therefore, we can instead perform an expansion of Eq. (2) in *T* and obtain the same expression in Eq. (4).

In the cases of both predominant latency and rapid transmission, within-host coalescence is restricted and prevents ‘short-sighted’ evolution, aiding long-term persistence in the population (Lythgoe et al., 2017). The reduction to a single-level coalescence described by the same function allows both fast-evolving RNA viruses and slow-evolving DNA viruses to be analysed using ARGs.

This is in contrast to HIV, which maintains a reservoir of latent CD4+ T-cells. Cycling through this reservoir maintains diversity that is transmitted much later (Doekes et al., 2017). With a large active compartment that quickly replicates, *ε* and *γ/R* are both non-negligible, leading to a dual-scale coalescent. Moreover, the coalescent genealogy is affected by treatment, which selects for lineages which may have been dormant for decades, thus warranting a structured or seed-bank coalescent (Dearlove and Wilson, 2013; Volz et al., 2017).

Therefore, although HIV undergoes short-sighted evolution by escaping our perturbative bound, the dormancy “seed-bank” retains a transmissible germline. While multiscale ARG inference that jointly models recombination and within/between-host coalescence remains a methodological challenge, we anticipate that variations on these perturbation methods could contribute to unpicking the dynamics of multiscale coalescents in HIV-1 and Ebola (Volz et al., 2017; Muñoz-Sánchez et al., 2026).

### Inferred recombination and its uncertainties

Our benchmarks suggest that the ARG approach and the ensemble statistical approach adopted by RDP5 present contrasting challenges. We distinguish between recombination events and breakpoints: an event is an instance of recombination between ancestral lineages, whereas a breakpoint is a genomic coordinate at which ancestry changes. RDP5 reports sequences inferred to be recombinant, identifies other sequences carrying evidence of the same event, and provides approximate start- and endpoints of the recombinant tract. Downstream analyses commonly convert these inferred events into fixed binary, count, or positional variables.

Yet, an observed sequence alignment may be statistically compatible with many combinations of historical events, inferred recombinant sequences, and breakpoint intervals. This ambiguity could arise from recurrent mutations, events that are not observable in the sampled genetic material, and variation in the local density of informative sites. Moreover, highly divergent regions with extensive homoplasy can obscure the mosaic patterns of recombination. These limitations are particularly relevant for analyses of EBV, where conserved regions, repeats, alignment masking, sequencing quality, and local diversity variation all influence how detectable a true recombination event is.

Our method accounts for uncertainty in the underlying recombination history by averaging over samples from the MCMC chain. Using SINGER’s output, we marginalise over event-level details, such as the branch, time, and coordinate at which a recombination node is placed. The posterior samples therefore indicate how strongly the sequence data support a change in ancestry within a genomic region.

In our benchmarks, TMRCA and genomic diversity were recovered accurately, motivating their use in our analysis. However, SINGER’s Bayesian framework regularises branch lengths through the neutral coalescent prior, which we expect to pull inferred node times towards the neutral expectation. Although sizeable deviations from the prior neutral expectation for TMRCA were seen in simulations, this shrinkage may attenuate the inferred deviations. Local TMRCAs therefore warrant cautious interpretation.

Using local tree topologies for downstream analysis also allows us to examine viral population structure by relating samples’ metadata to their placement in the phylogeny. We anticipate that this representation could help distinguish an old ancestry transition inherited by a large clade from several recent changes, extending the analysis beyond the presence of a recombinant tract in an alignment. Furthermore, local tree spans are inferred rather than specified by gene boundaries, providing a functionally agnostic alternative to gene trees and whole-genome trees.

### ARGs provide a description of evolutionary modularity

Human demography and the immune system are tightly coupled by shared ancestral migratory history, environmental challenges, and pathogen-driven HLA diversification (Arrieta-Bolaños et al., 2023; Liston et al., 2016). Since EBV transmission is highly assortative, demographic structure can generate geographic associations that resemble locus-specific selection. Local genealogies enable these associations to be compared across the genome, with the ARG providing a demographic null expectation shared across all loci. The absence of geographic association at *BFLF1*, an early lytic gene with no known immune pressure, supports this idea. In immunogenic protein-coding genes, selective pressure may preserve particular allelic combinations despite homologous recombination and therefore contribute to the structure EBV selection (Briercheck et al., 2024), observed as the slow LD decay over physical distances.

The deep type 1/2 axis in *BZLF2* with low within-type diversification relative to between-type diversification is consistent with its molecular function: gp42 engages HLA class II molecules through a constrained binding surface (Spriggs et al., 1996). gp42 also functions as an entry tropism switch, which further delineates its constraint landscape from gL/gH.

The intermediate diversification timescale of *BKRF2* (gL) and high variance within geographical clusters indicates continuous diversifying pressure that generates allelic novelty faster than homogenising drift, in line with the accelerated evolution described by Santpere et al. (2014).

gH diversification must simultaneously maintain binding interfaces with gL, gp42, the gB fusogen, and the KGD motif that binds epithelial integrins (Sathiyamoorthy et al., 2016), supporting the observation that gH’s within-cluster divergence is an order of magnitude smaller than gL’s.

*BKRF2* -*BXLF2* (gL-gH) exhibit a mean RF distance of 162.7, compared to 137.1 for gp42-gL and 170.7 for gp42-gH), yet Figure 4B shows that LD between gH and gL is elevated relative to their LD with gp42. Thus, the stronger allelic association between gH and GL does not correspond to the greatest similarity in their inferred tree topologies. A potential explanation is epistatic co-selection maintaining compatible binding interfaces, with geography superimposed: the two constraints are intertwined.

An alternative hypothesis to long-range epistasis is population history. Human herpesvirus 8, also a gammaherpesvirus, exhibits higher diversity at the K15 locus than surrounding genomic regions. It has been hypothesised that this gene entered the KSHV genome via introgression from a deeply divergent, probably extinct, KSHV-like virus (Hayward, 1999). A similar history in EBV could produce deep divergences at individual loci without requiring epistatic selection to explain this divergence. Further work can distinguish whether the observed allelic associations in EBV can be explained by the inheritance of divergent ancestry, or require selection to maintain high-fitness combinations despite recombination.

In either case, these findings have potential implications for vaccine design: candidates targeting individual glycoproteins implicitly assume that target antigenic variation co-varies with other entry components. Our ARG analysis indicates that recombination combinatorially expands the antigenic space; instead, structural conservation at inter-protein interfaces may provide a more stable antigenic target, as empirically demonstrated by gL/gH/gp42 complex-targeting antibodies (Bu et al., 2019).

## Conclusions

These examples corroborate our hypothesis that recombination accommodates evolutionarily modular assemblies that answer to different constraints. The ARG captures a richer subspace of the selection landscape than locus-specific or global methods, and unifies phylogenetic explanations from Zanella et al. (2019) and Santpere et al. (2014). Nevertheless, our characterisation of gH/gL’s variation would be strengthened by HLA-typing and a more structured approach to geographical/demographic labelling (Briercheck et al., 2024; Lynn-Green et al., 2023).

A practical implication is that disease-association studies should prioritise geographically matched case-control designs, as in (Wong et al., 2024; Ji et al., 2025; Zhang et al., 2025). Pairing EBV phylogenetic information with host genotype/HLA data could reveal host-pathogen dynamics relevant to cooperative EBV persistence (Styles et al., 2018), EBV vaccines that provide sterilising immunity against infection (Soldan et al., 2022; Giovannoni et al., 2025), and epigenetic pathogenesis (Li et al., 2018). Ultimately, our ARG-informed framework provides a new population genetics lens that integrates over topological uncertainty. Ongoing efforts to increase the quality and quantity of EBV sequences (Xie et al., 2024) will help untangle the complex interplay of viral latency, immune evasion, and oncogenesis.

## Methods

### Data and pre-processing

All analyses were performed on a whole-genome multiple-sequence alignment of 147 Epstein-Barr virus genomes from Wegner (2017); Correia et al. (2018), and joined with per-sequence metadata (type 1/2 EBV, geographic area, cancer/normal status, and associated disease) Gene and repeat annotations were converted to alignment coordinates, and repetitive regions were excluded: these intervals were merged into a minimal set of disjoint intervals, allowing all downstream analyses to exclude these regions.

### Estimation of population genetic parameters

Writing the number of segregating sites *S* over *L* callable loci, and 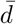 for the average number of pairwise differences computed over all pairs with gaps and ambiguity codes excluded pairwise, we computed the following quantities for an assumed per-site, per-generation mutation rate *µ*:

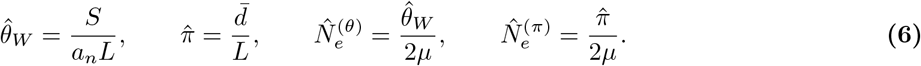

where *a*_*n*_ is the *n*th harmonic number. We use 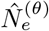 to initialise SINGER. Given the coalescent model, a ‘generation’ in the ARG framework corresponds to the mean serial interval between transmission events, instead of a viral replication cycle.

Assuming a standard dsDNA viral substitution rate of ≈ 1 *×* 10^−8^ site^−1^year^−1^ and a 10-15 year interval, compounded by a mutational penalty driven by lytic replication, we set *m* = 2.5 *×* 10^−7^ generation^−1^.

RAxML (Stamatakis, 2014) was used as a baseline topology for recombination rate inference, which was passed to ClonalFrameML (Didelot and Wilson, 2015) to estimate relative importation densities per site, *d*(*i*) and global *R/θ*. We scale *d*(*i*) by the latter to find 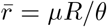, and find absolute local recombination rates 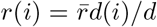. *r*(*i*) was averaged into non-overlapping 1kb windows. To prevent MCMC chain collapse, windows with zero inferred coverage were floored to 1 % of 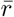.

## SINGER

We inferred 150 ARGs, with the following parameters: Convergence was confirmed from the sampled tree sequences, following methods from Deng et al. (2025) (Figure 7). First, we computed the number of incompatible sites (sites with more than two mutations, or exactly two mutations, neither of which sits on a root branch). Second, we fit a diversity-fit statistic measuring the agreement between the mutational and genealogical scales,

**Figure 7.**
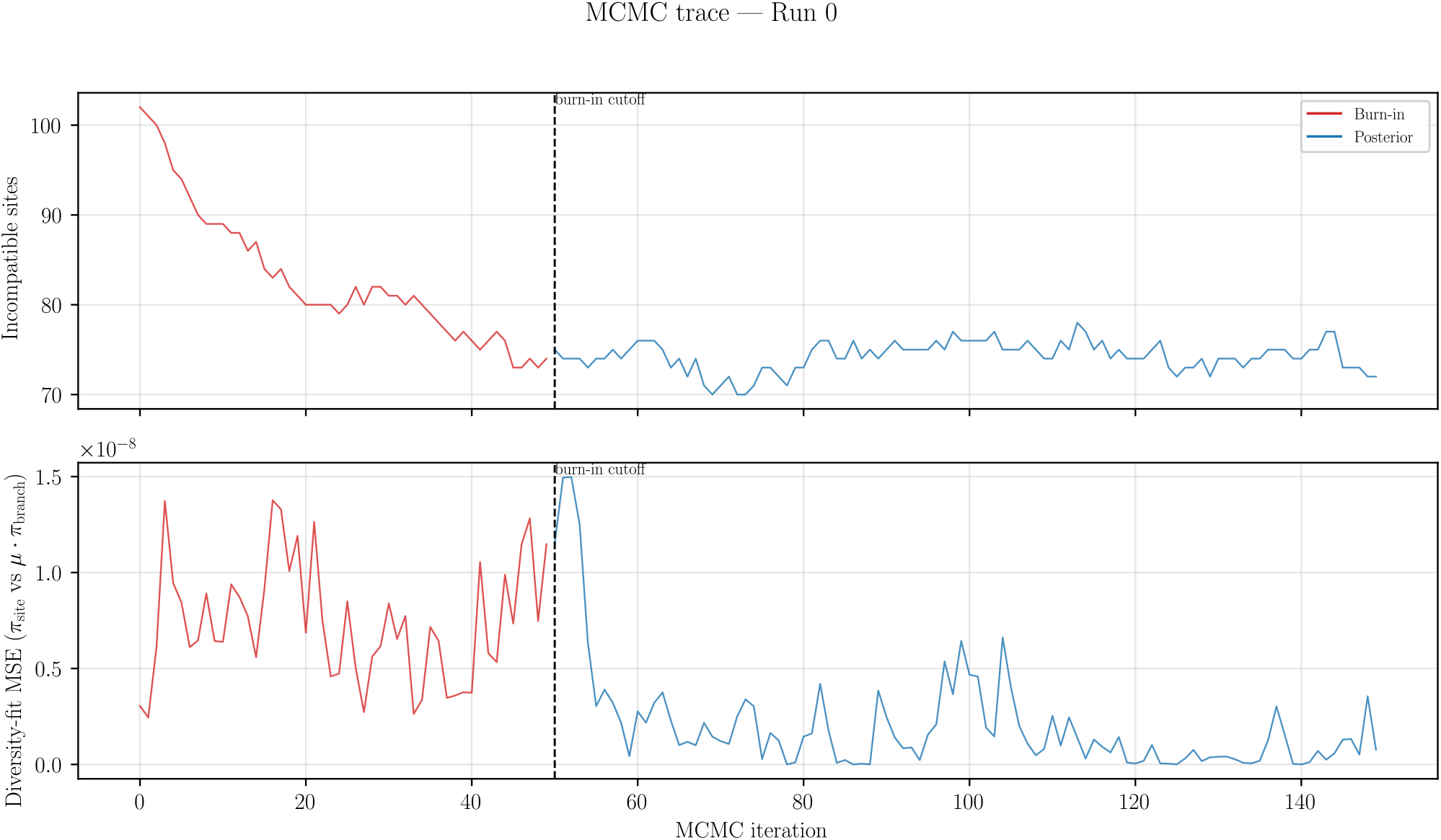
Number of incompatible sites in the ARG and *π*_site_ against *µ* · *π*_branch_ across the MCMC iterations, with the dotted line indicating the burn-in cutoff.

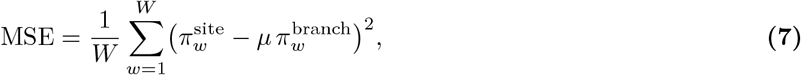

evaluated over 1 Mbp windows, where *π*^site^ and *π*^branch^ are the site- and branch-mode nucleotide diversities.

### Simulations

Coalescent-with-recombination genealogical histories were simulated with msprime under a haploid Wright– Fisher scaled neutral model with parameters *N*_*e*_, *ρ*, and *L*, and the same number of sample haplotypes. For each *r/m* ratio (0.02, 0.05, 0.1, 0.2, or 0.5), 10 independent calibration and 20 independent test datasets were generated. An interval was labelled positive for recombination if the true msprime ARG contained a recom-bination event there. SINGER scored each interval as the fraction of 150 retained posterior ARGs containing a parental lineage switch, with ROC analysis sweeping across the consensus threshold. RDP5’s breakpoint endpoints were mapped to the same intervals, and using the reported p-value rankings to sweep across reported breakpoint intervals.

### Analyses

#### Gene clustering

We bin the sequences at 100bp resolution. For each posterior ARG sample, the Robinson-Foulds distance was computed between all pairs of distinct marginal trees, yielding a symmetric matrix *R*^(*k*)^, and the posterior mean 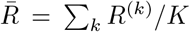 is reported. Average per-ARG pairwise divergence for each gene interval was given by

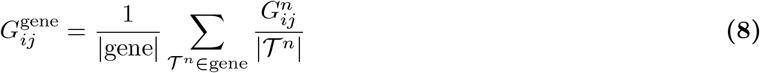

The distance matrices underwent hierarchical UPGMA clustering for *k* ∈ [2, 6]: averaging over the posterior distribution smooths stochastic rate variation across lineages, meaning the distance matrices more closely approximate ultrametricity than single-tree or sequence-based methods do. Optimal *k* was selected using silhouette scoring, and outliers were checked manually and removed. Metadata-cluster association was tested by Fisher’s exact test for two-cluster genes and otherwise by label-shuffling *χ*^2^ permutation tests (*n* = 10000).

## Supporting information

Raw data

## Data availability

All code and scripts are available at https://github.com/leungchristopher/viralarg.The haploid version of SINGER is available at https://github.com/leungchristopher/SINGER, with test accelerated versions in branches. Upon publication, a pull request will be made to incorporate these features into the package at https://github.com/popgenmethods/SINGER.

## Acknowledgements

We thank Richard Durbin for useful discussions.

## Author contributions

C.H.C.L., all analyses, software development, and drafting of the manuscript; C.J.H. and A.S., supervision and manuscript revision.

## Supplementary Information

### The application of perturbation theory to genetic models

Perturbation theory is a class of mathematical methods used to find approximate solutions to problems that can be divided into soluble and perturbative parts. When the genetic model is governed by a linear operator with a biologically small parameter, perturbative methods give an asymptotic approximation of the genealogy, and have found applications in problems such as selection-mutation models (Stephens et al., 2005).

The initial distribution ***π***_0_ = (0, 0, 0, 1) is the equilibrium of the unperturbed chain, occupied with probability 1 − *O*(*ε*). For finite *T* the matrix exponential is entire in *ε*, so the series converges for all *ε* (Kato, 1995). Yet small *ε* gives us accuracy upon truncation.

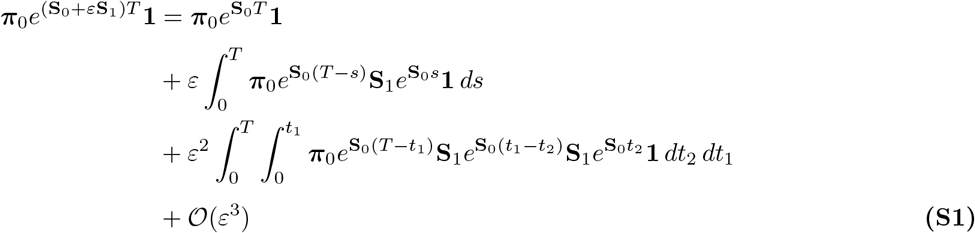

Two intermediate quantities recur. Let 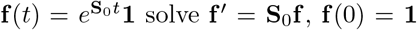 the survival probability of the unperturbed chain from each state. States 2, 3 and 4 cannot coalesce without a reactivation, so *f*_2_ = *f*_3_ = *f*_4_ = 1, while state 1 decays into them,

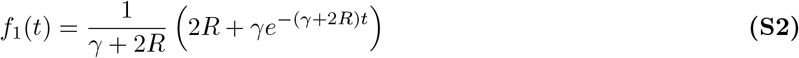

giving 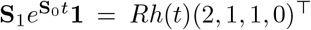 with *h*(*t*) = *f*_1_(*t*) − 1 ≤ 0. Second, state 4 is absorbing under **S**_0_, so 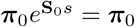 for all *s* and

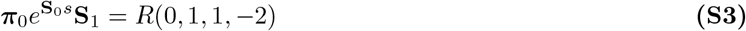

The first reactivation is thus equally likely to hit either of the two lineages being considered, and its timing is irrelevant: the pair waits in state 4 until it occurs.

To solve the second-order term, with *τ* = *t*_1_−*t*_2_ and 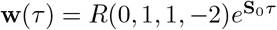, the Kolmogorov forward equation **w**^*′*^ = **wS**_0_ gives *w*_1_ ≡ 0 – state 1 is unreachable from a singly lytic state without a further reactivation – and

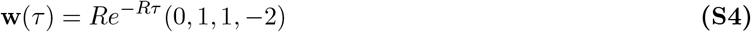

The exponential term represents the required coincidence of two lytic lineages. Contracting Eq. (S4) with *Rh*(*t*_2_)(2, 1, 1, 0)^⊤^ leaves 2*R*^2^*h*(*t*_2_)*e*^−*Rτ*^, and

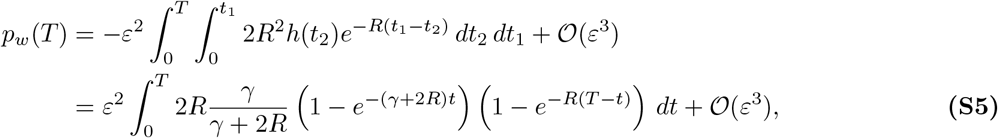

Letting *a* = *γ* + 2*R*,

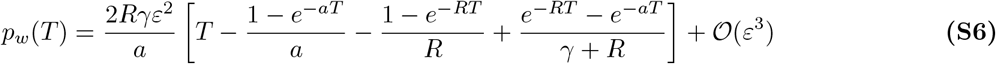

Since the spatial separation of different lineages means that *γ* ≪ 2*R*, the prefactor can be truncated to give 2*Rγ/a* = *γ*(1 − *γ/*2*R* + …). A long co-infection removes the exponentials, which are boundary layers of width *R*^−1^ at each end of the interval, leaving the bracket as *T* − 3*/*(2*R*). Hence

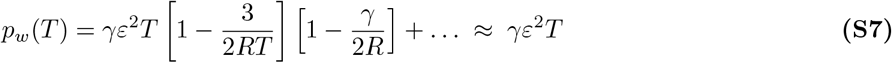

We can treat pairs as coalescing independently, as the probabilities are individually *O*(10^−8^), making them effectively disjoint.

If *ε* is not small, and *γ* is not negligible relative to *R*, the series converges but the low-order truncation is not valid. This represents the regime of chronic, but highly active infections such as HIV and hepatitis C (Vrancken et al., 2015; Riaz et al., 2022), in which competing lineages persist within a single host and within-host coalescence cannot be neglected.

### Computational optimisations to SINGER

First, SINGER was modified to support a haploid population genetics model, which incidentally also doubles the computational limit on the number of sequences. Second, full support for 1kb windowed recombination rates was added to SINGER. When a recombination map is passed, it builds a matching mutation map with the same interval boundaries in `compute_mut_map()`. This is then converted into a cumulative, integrated-rate representation in `Rate_map.cpp. Sampler.cpp` then creates an internal reference grid, onto which the recombination rate map is projected by integration: 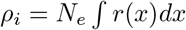 over each interval block in `ARG.cpp`.

Performance-oriented data structures were also implemented. We replace `std::map, std::set` containers with flat sorted arrays for the coalescent calculator, and fix problems with dereferencing and invalidated iterators. This should prevent edge-case proposals from crashing the MCMC chain.

